## Supplementary Materials for "Dynamical modeling of the H3K27 epigenetic landscape in mouse embryonic stem cells"

**Table S1: Tested combinations of $r_{13}$ and $r_{23}$.** Over all the tested cases, only $r_{13} = r_{23} = 3$ leads to a satisfying fit of the experimental profiles of H3K27 modifications around PcG-target genes (Fig.S3). Failure of the combination $r_{13} = r_{23} = 4$ is illustrated in Figure S4.

| $r_{13}$ | $r_{23}$ |
|---|---|
| 9 | 10,9,8 |
| 6 | 7,6,5 |
| 4 | 5,4,3 |
| 3 | 4,3,2 |
| 2 | 3,2,1 |
| 1 | 2,1,3 |

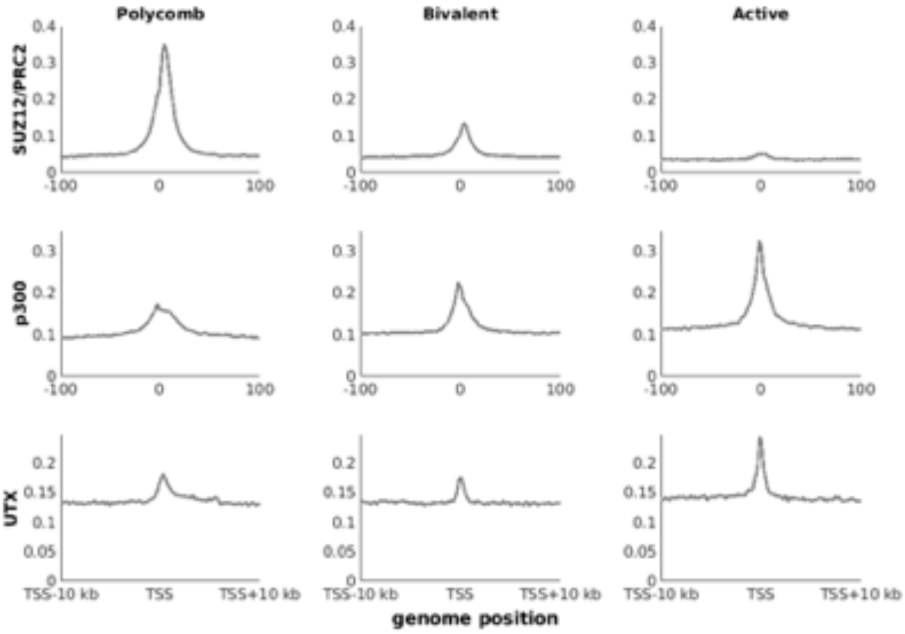


**Fig.S1**: **HMEs profiles.** Average Chip-seq densities (normalized) of SUZ12, p300 and UTX of PcG-target, bivalent and active genes around the TSS in WT condition.

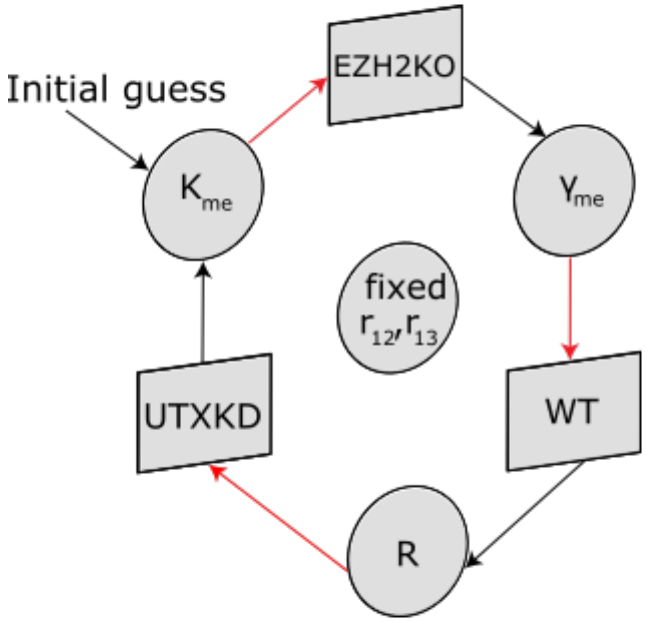


**Fig.S2: Iterative inference strategy.** For fixed values of $r_{12}$ and $r_{23}$, an initial guess for $k_{me}$ is used to initialize an iterative inference cycle where a parameter inferred at one step feeds (red arrows) the next inference step based on various datasets (black arrows): EZH2KO data to infer $\gamma_{me}$, WT for $R$ and UTXKD for $k_{me}$ (see main text).

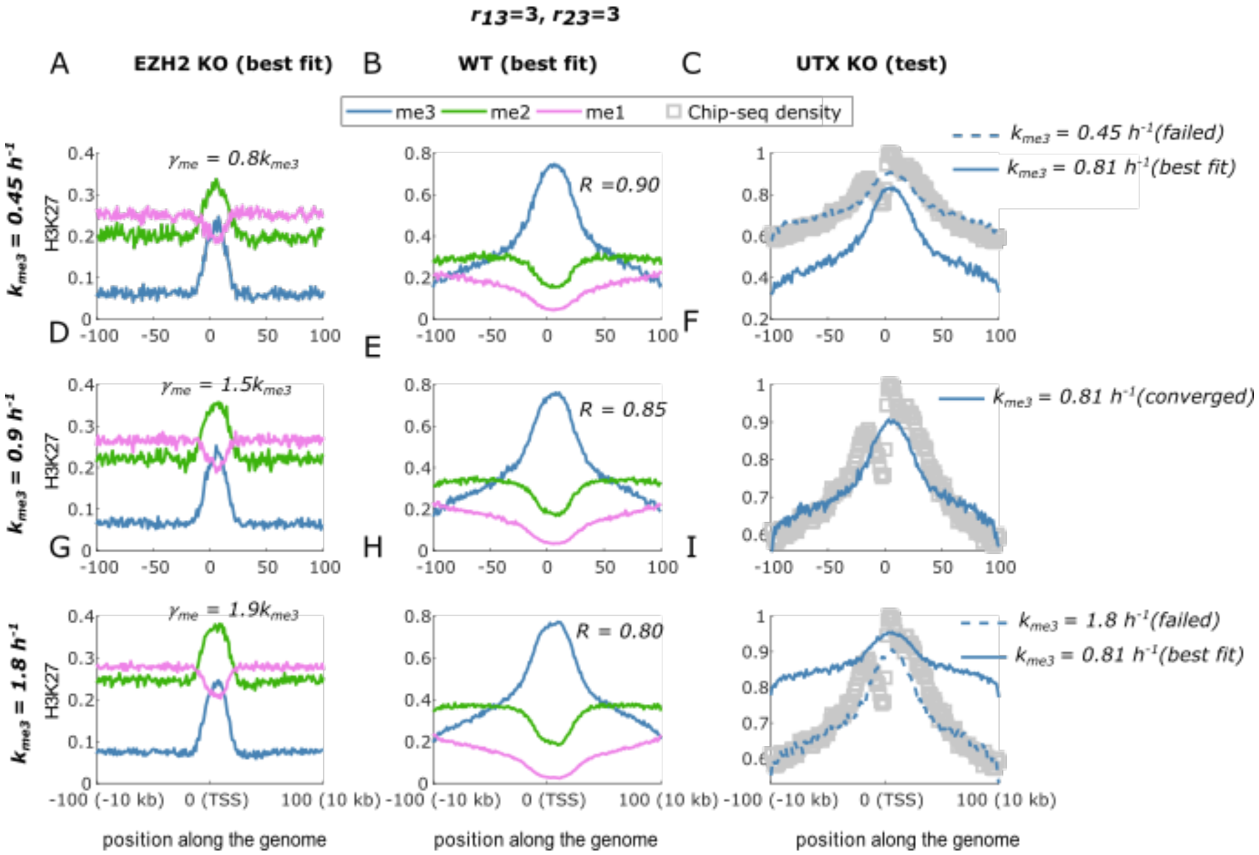


**Fig.S3**: **Parameter inference for parameter** $r_{13} = 3$**,** $r_{23} = 3$. The steps for fixing $R, k_{me3}$ and $\gamma_{me}$ are illustrated. (A,D,G) H3K27 methylation are best fitted to EZH2 KO experimental profile by fixing $\gamma_{me}$for a particular $k_{me3}$. (A,D,G) Then, H3K27 methylation are best fitted to WT experimental profile by fixing $R$ for a fixed pair $k_{me3}, \gamma_{me}$. (C,F,I) Finally the fixed parameters $k_{me3}, \gamma_{me}, R$ are tested if the H3K27me3 profile fits experimental density.

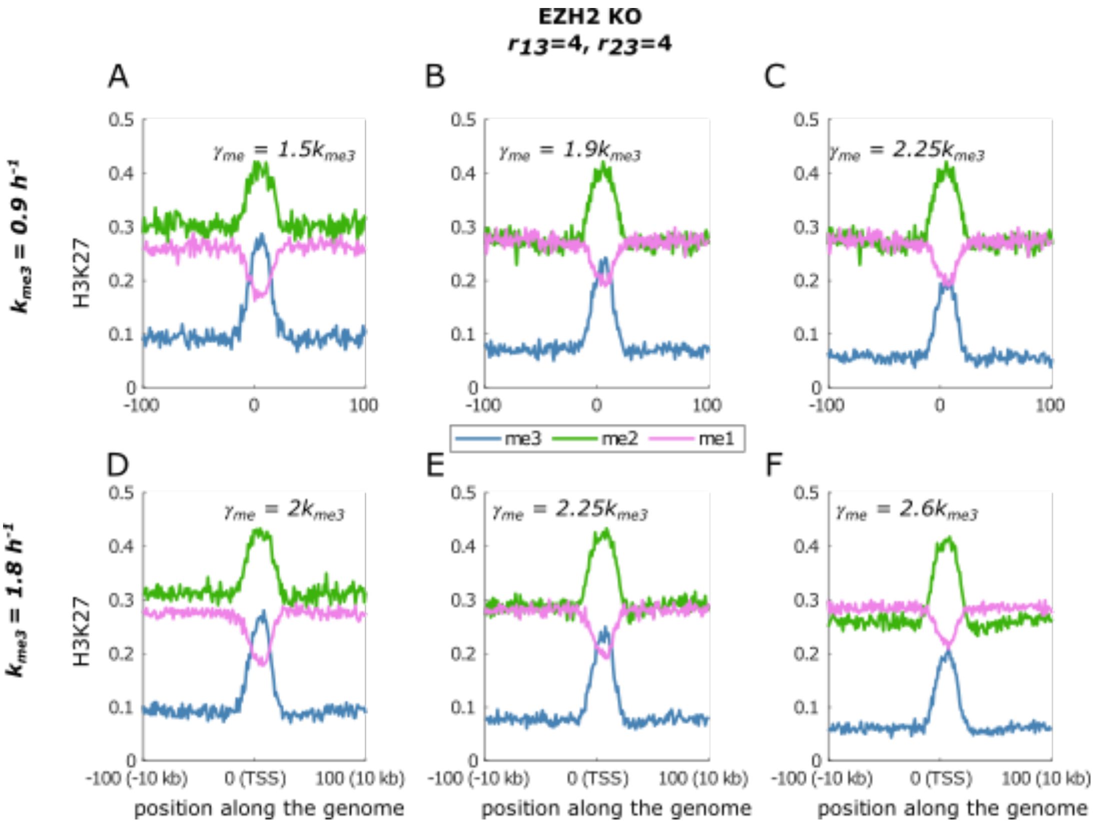


**Fig.S4: Parameter inference for parameters** $r_{13} = 4$**,** $r_{23} = 4$. With these parameters, simulated H3K27 methylation profiles of EZH2 KO never capture the experimental methylation valency at promoters (Figure 2F of the main text). Top panel is for $k_{me3} = 0.9\ h^{-1}$and explores $\gamma_{me}$ to find a suitable $k_{me3}, \gamma_{me}$pair to qualitatively capture methylation valency of EZH2KO. Bottom panel explores $\gamma_{me}$for $k_{me3} = 1.8\ h^{-1}$.

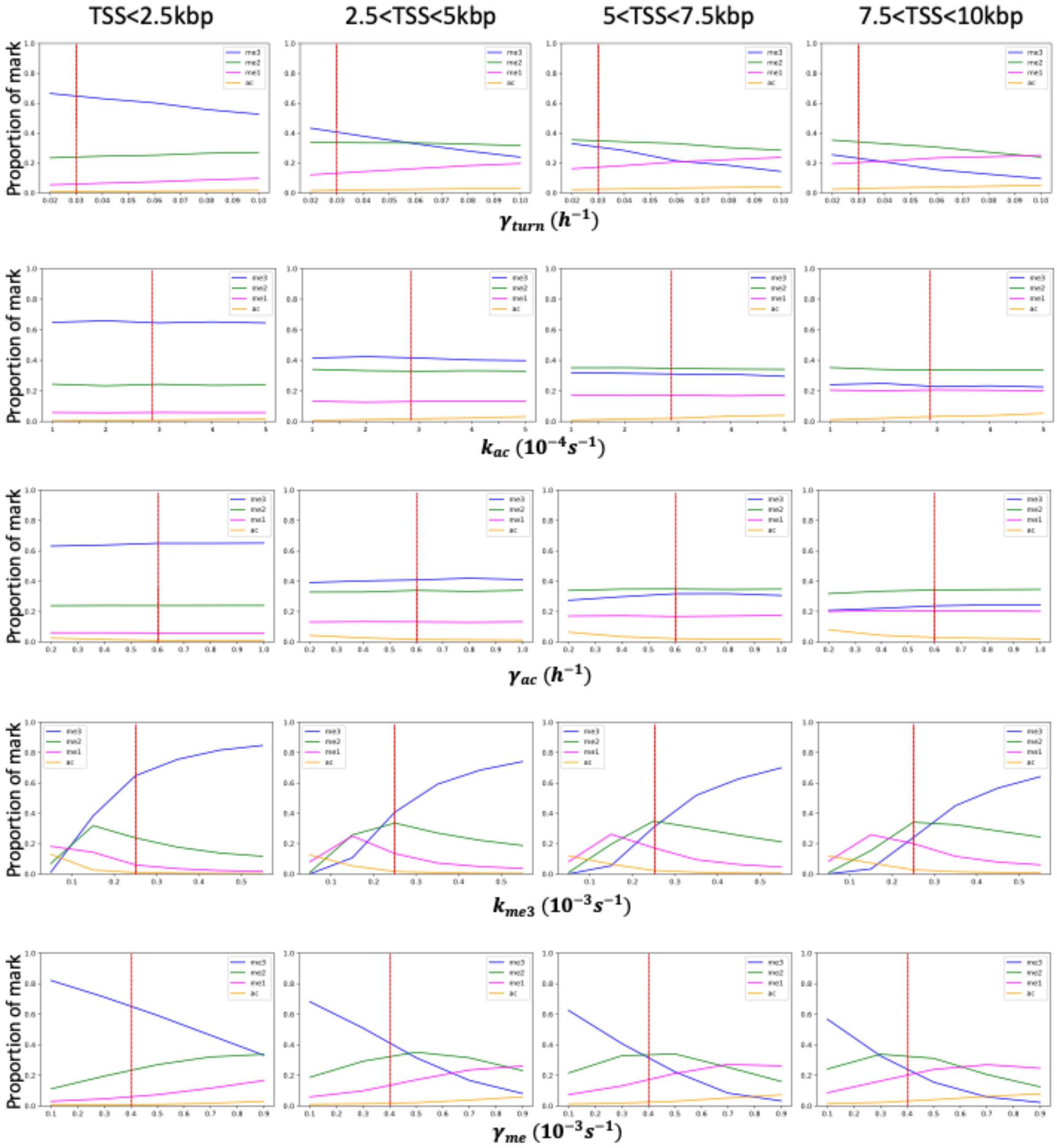


**Fig.S5: Systematic variation of model parameters.** Average predicted proportion of a given mark around PcG-target genes as a function of the different model parameters, all other parameters fixed to WT values (red dotted lines). Panels from left to right correspond to regions close or far from TSS. For $k_{me3}$, we also varied $k_{me1}$ and $k_{me2}$ by keeping $r_{13}$ and $r_{23}$ constant to WT values (see Table 1 of the main text).

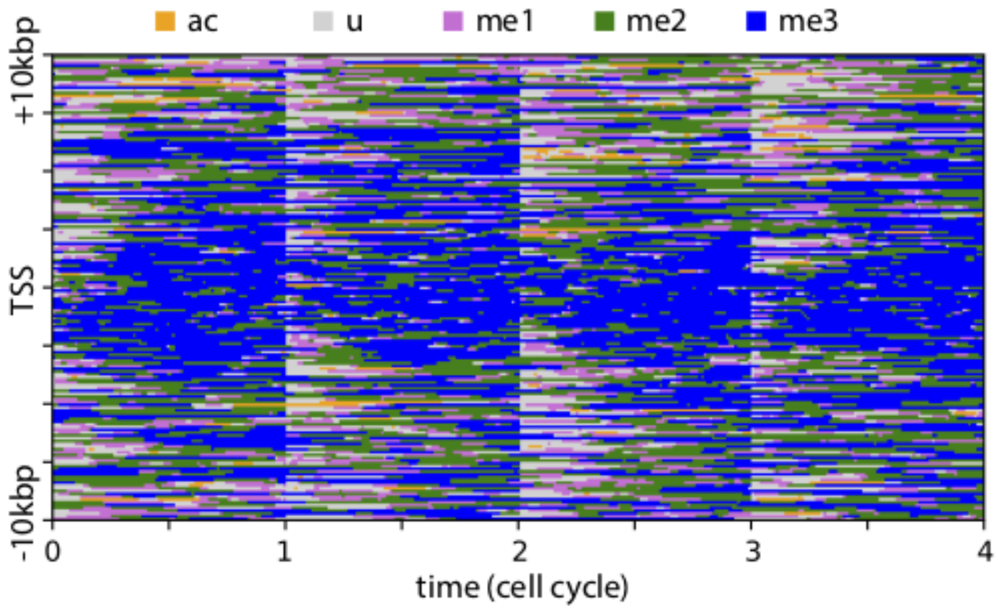


**Fig.S6**: **Stochastic dynamics**. Kymograph representing a typical simulation trajectory at periodic steady-state around PcG-target genes with WT parameters obtained with the Gillespie algorithm. The local epigenetic state fluctuates stochastically following the kinetic rates given in the text.

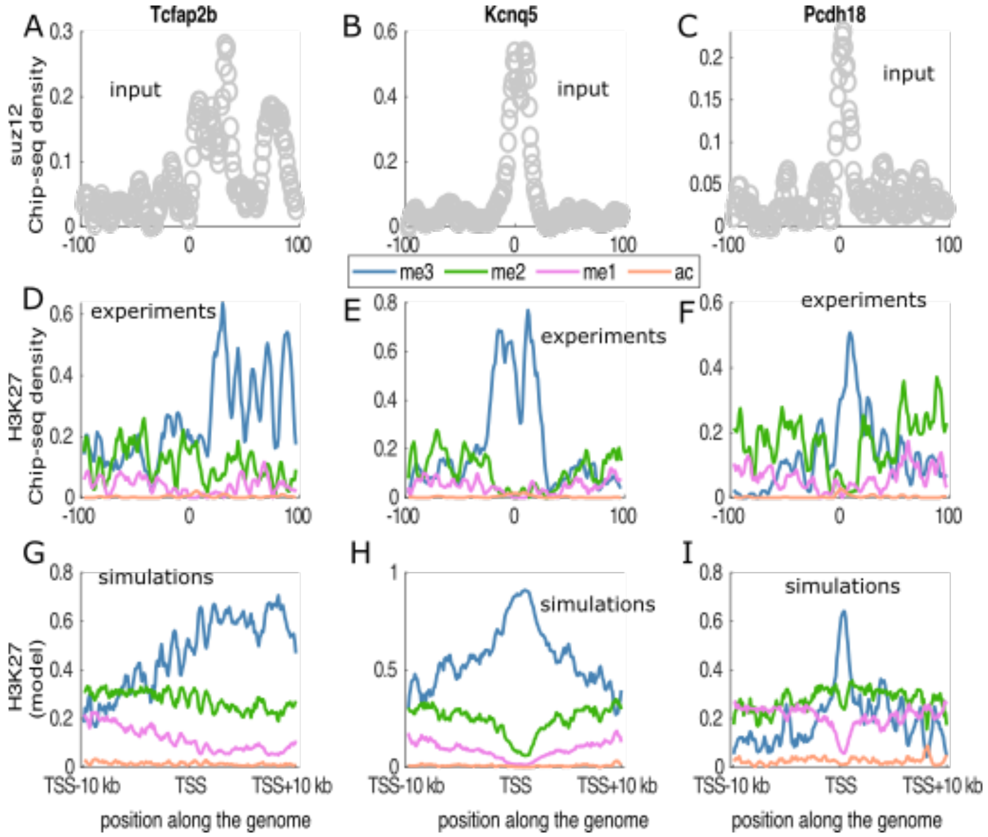


**Fig.S7**: **Predictions of single PcG-target genes**. Predicting the H3K27 modification landscape at single genes in PcG-target domains. Profiles of gene *Tcfap2b* (left column), *Kcnq5* (middle column), and *Pcdh18* (right column). (First row) Input SUZ12 occupancies of three specific genes. (Second row) Chip-seq H3K27 methylation and acetylation for corresponding genes. (Third row) Simulated H3K27 modification profile of the respective genes.

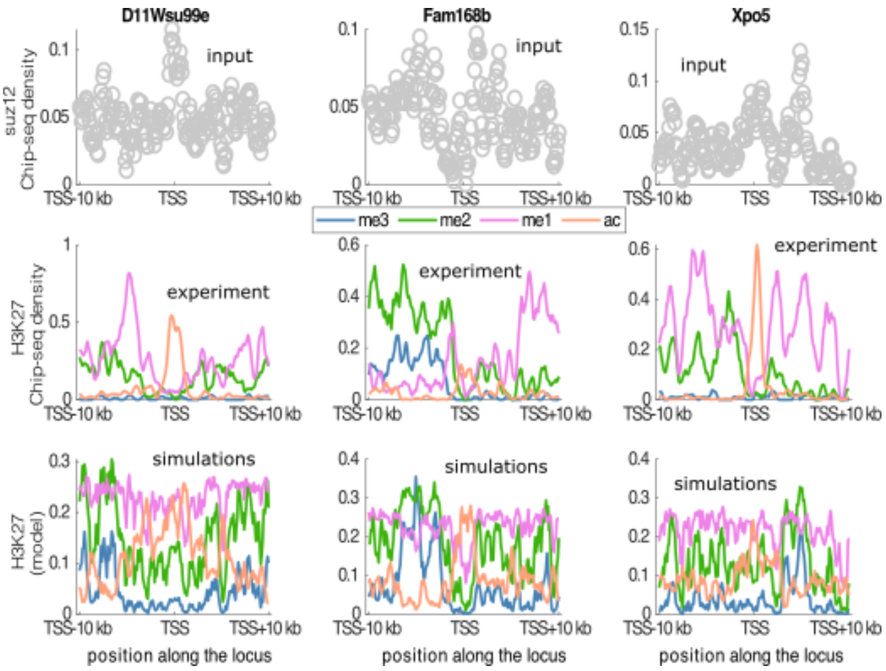


**Fig.S8: Predictions of single active genes.** Predicting the H3K27 modification landscape at single genes of active domainProfiles of gene *D11Wsu99e* (left), *Fam168b* (middle) and *Xpo5* (right). (First row) Input SUZ12 occupancies of three specific genes. (Second row) Chip-seq H3K27 methylation and acetylation for corresponding genes. (Third row) Simulated H3K27 modification profile of the respective genes.

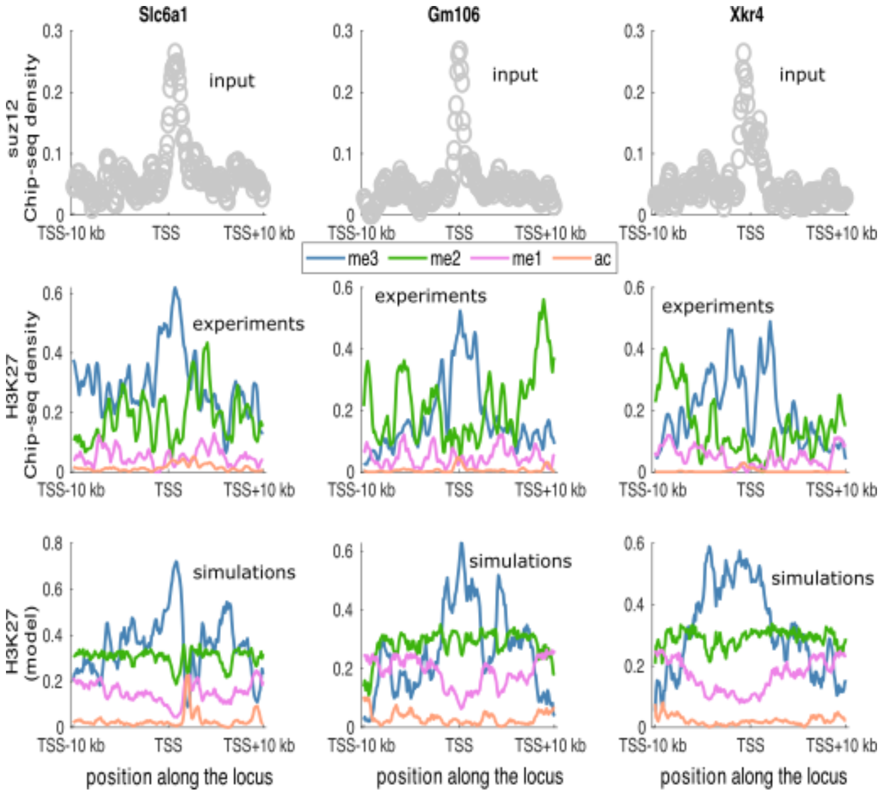


**Fig.S9**: **Predictions of single bivalent genes.** Predicting the H3K27 modification landscape at single genes of bivalent domain. Profiles of gene *Scl6a1* (left), *Gm106* (middle) and *Xkr4* (right). (First row) Input SUZ12 occupancies of three specific genes. (Second row) Chip-seq H3K27 methylation and acetylation for corresponding genes. (Third row) Simulated H3K27 modification profile of the respective genes.

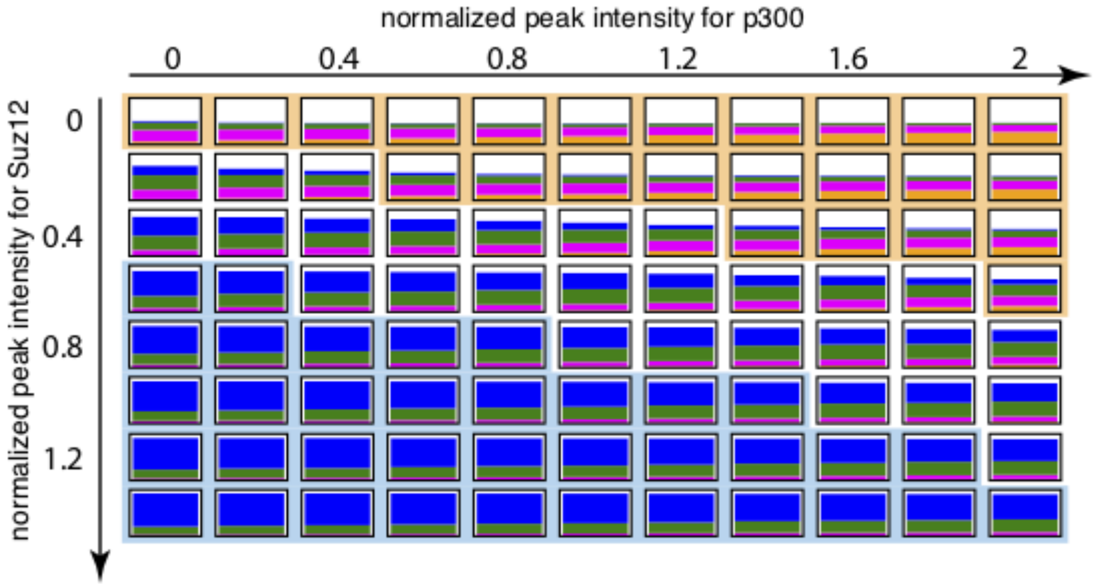


**Fig.S10: Competition between HMEs**. We varied the strengths of recruitment of p300 (x-axis) or Suz12/PRC2 (y-axis) around TSS for WT parameters. For each condition, we computed the average proportion of each H3K27 mark in a $\pm 2.5$kbp window around TSS. The corresponding stacked bar charts are given in the subplot (blue: me3, green: me2, magenta: me1, orange: ac, white: u). This allows us to define qualitatively two regions depending on the relative methylation valency: a PcG-target-like region (blue area) with me3>>me2>me1 and a active-like region (orange area) with me1>me2>>me3.

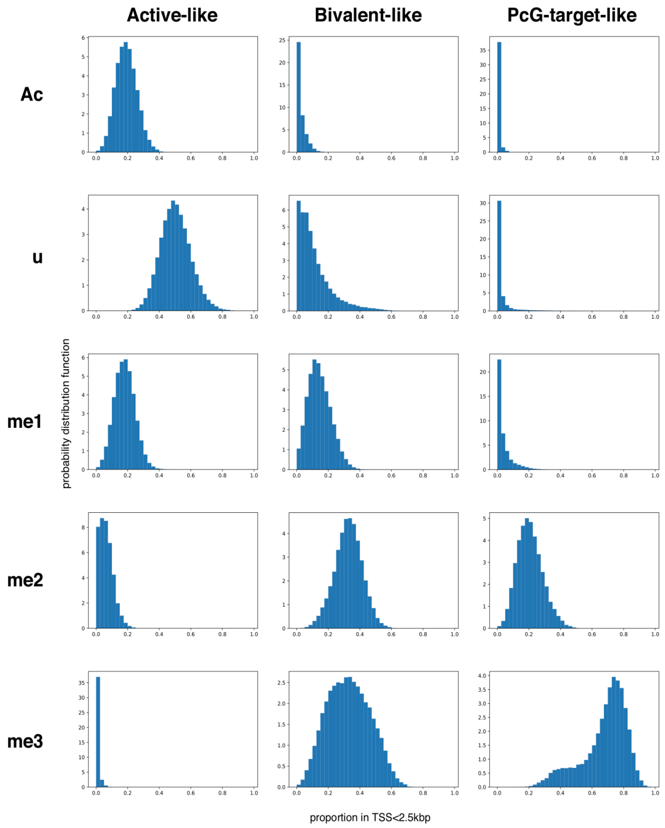


**Fig.S11**: **Distribution of H3K27 states.** Probability distribution functions for the proportion of a H3K27 state inside the region TSS$\pm$2.5kbp in a population of unsynchronized cells for three values of HME recruitment strengths (Fig.5A of the main text), one in the Active-like region ($(\alpha;\beta) \approx (0;1)$), one with a bivalent-like inputs ($(\alpha;\beta) \approx (0.4;0.5)$) and one in the PcG-target-like region ($(\alpha;\beta) \approx (1;0.3)$).

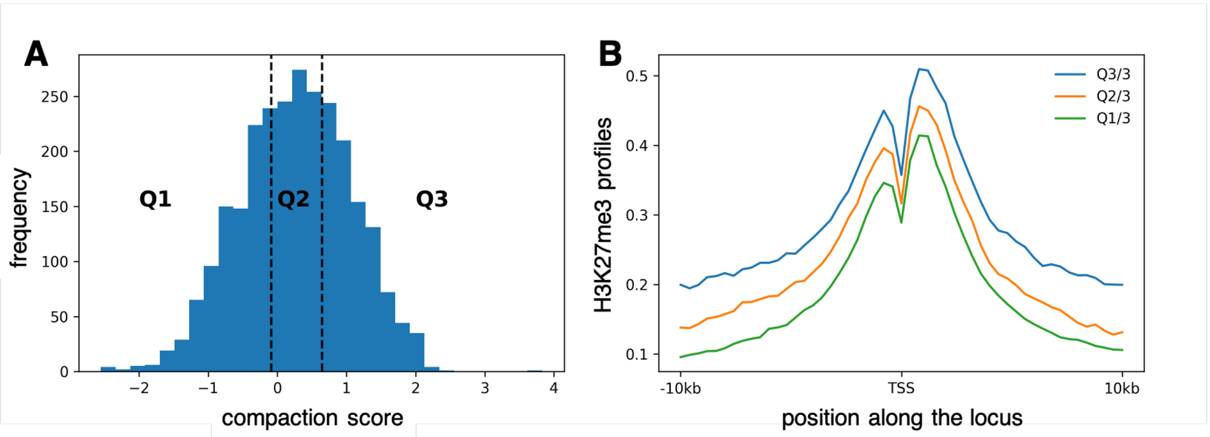


**Fig.S12: Relation between compaction and H3K27me3 profiles.** For each PcG-target gene, we estimated a compaction score that translates the density of 3D contacts around this gene. More precisely, we took the Hi-C data of mESCs at 10kbp resolution from (Bonev et al, Cell, 171: 557–572.e24, 2017) that we distance-normalized (the Hi-C value of each bin *(i,j)* is normalized by the average contact frequency at genomic distance |*j-i*|) to obtain the so-called observed-over-expected contact matrix *OE*. For a gene *g* with a TSS at position $i_g$ along the genome, we define its compaction score as the log2 of the median value of the OE matrix in a $\pm 100kbp$ window around the TSS: $log_2[median\{OE((i_g - 100kbp):(i_g + 100kbp);(i_g - 100kbp):(i_g + 100kbp))\}]$. The distribution of compaction scores in the ensemble of PcG-target genes is given in panel (A). We divided this ensemble into three subgroups of the same size: Q1 with low compaction scores, Q2 with intermediate and Q3 with high scores (A). Panel (B) shows the average H3K27me3 profiles around TSS for each subgroup (computed as the other average H3K27 profiles in the main text). The more compact the gene is the more extended the profile is.

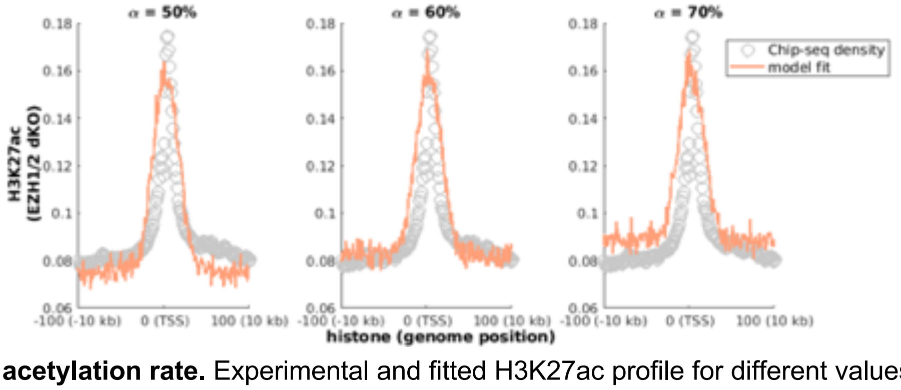


**Fig.S13**: **Fit of acetylation rate.** Experimental and fitted H3K27ac profile for different values of $\alpha$. The best fit $\alpha = 0.6$ is picked for which $k_{ac} = 1.03\ h^{-1}$.
